# Tool recommender system in Galaxy using deep learning

**DOI:** 10.1101/838599

**Authors:** Anup Kumar, Björn Grüning, Rolf Backofen

## Abstract

Galaxy is a web-based and open-source scientific data-processing platform. Researchers compose pipelines in Galaxy to analyse scientific data. These pipelines, also known as workflows, can be complex and difficult to create from thousands of tools, especially for researchers new to Galaxy. To make creating workflows easier, faster and less error-prone, a predictive system is developed to recommend tools facilitating further analysis. A model is created to recommend tools by analysing workflows, composed by researchers on the European Galaxy server, using a deep learning approach. The higher-order dependencies in workflows, represented as directed acyclic graphs, are learned by training a gated recurrent units (GRU) neural network, a variant of a recurrent neural network (RNN). The weights of tools used in the neural network training are derived from their usage frequencies over a period of time. The hyper-parameters of the neural network are optimised using Bayesian optimisation. An accuracy of 97% in predicting tools is achieved by the model for precision@1, precision@2 and precision@3 metrics. It is accessed by a Galaxy API to recommend tools in real-time. Multiple user interface (UI) integrations on the server communicate with this API to apprise researchers of these recommended tools interactively.

## Introduction

Life sciences depend increasingly on high-throughput data (HTD), turning them into data science to a large extent. However, raw HTD does not have much value on its own without proper analysis and interpretation of the data. To simplify the data analysis process and to ensure a reproducible analysis, several workflow engines have emerged ((20); (14);(2)). The main idea for workflow systems is based on the observation that any computational analysis of HTD encompasses multiple steps such as quality control, preprocessing, quantification and statistical analysis to transform raw data into scientific results. Collectively, these steps form a workflow where each step performs a definite transformation of the data, which can be performed using standardised tools. Using workflow for the analysis is simple and convenient and has several advantages. First, it is easy to replace individual tools by a newer version or to assess the influence of the associated step on the final result. Second, a workflow can be saved, shared and used multiple times, which ensures reproducible research. Therefore, workflows are becoming essential in the analysis of scientific data and there are multiple platforms where researchers can create workflows for their analyses. However, a critical question is how to assess whether a generated workflow is State-of-Art or even valid at all. To give a concrete example, one can use several real-valued input vectors (such as fluorescence-based measurement stemming from arrays), transform them into integer-based values in the first step and combine it with a tool that uses a count-based statistics (such as negative binomial distribution as used in DeSeq2) to determine values that show high differential behaviour. While this workflow would run on a workflow system without problems and even produce some results, the generated results are not valid because of the wrong statistical model.

Galaxy is a open-source data processing platform which enables researchers create and store their workflows for multiple scientific analyses ((1)). A workflow in Galaxy is a directed acyclic graph and consists of one or many tool sequences to analyse scientific data such as DNA and RNA sequences. A tool consumes one or more data files as input and produces one or more data files as output and has a defined number of data types for these input and output files. In workflows, the tools are connected one after another following a constraint that the adjacent tools must have compatible data types. In other words, the data types of output files of a tool should match the data types of input files of the following tool. Galaxy has thousands of accessible tools and acquiring familiarity and constructing workflows with these tools can be a complex and time-consuming task, especially for researchers new to Galaxy. To assist them in creating workflows and making them aware of the possible tools for further analyses, a recommender system is devised. The benefits of having such a system are manifold. First, it will avoid the loss of time spent in creating erroneous or less optimal workflows by choosing tools which may be wrong and thereby making researchers more efficient. Second, it will help them bypass the step of searching for tools separately which shows potential to further reduce the time spent in creating workflows and increase the accessibility of tools. Third, it will promote tools having higher usage frequencies in the past to the top of the recommendations and downgrade those having lower usage frequencies to the bottom of the recommendations. It is achieved by assigning weights to tools which are derived from their usage frequencies over a period of time. Finally, it can also be used to promote the newly added tools in Galaxy by showing them alongside the recommended tools predicted using the deep learning approach.

### A. Recommender systems

The objective of having recommender systems in fields such as scientific literature search, online shopping, travel bookings, media-service providers and many other fields is to let people discover suitable, interesting and the newly-released products. These recommended products are recognised based on the usage and purchasing patterns of people in the past. In the field of scientific literature search, the exponential increase in the number of published papers necessitates having a recommender system to help scientists explore relevant and recent papers quickly ((3)). Companies such as Amazon and Netflix have appropriately used recommender systems to learn preferences of their respective customers in selecting products such as their favourite books or movies and to propose a few products out of a large store. It becomes faster for their customers to sift through a few recommended products to find the most suitable ones rather than looking in their complete stores. By enabling their customers discover reasonable and customised products, the recommender systems have helped these companies grow as organisations ((32); (31)). The successful implementations of recommender systems by many organisations across the world working in diverse areas to assess the needs of their customers in choosing relevant items and to propose the most useful ones motivated us to create a tool recommender system in Galaxy.

### B. Related work

To simplify creating workflows for scientific analyses a few approaches have been proposed which suggest alternative tools and workflows. ((25)) makes use of EDAM and semantic annotations of tools to compose workflows automatically for mass-spectrometry based proteomics. The annotations include the names, functionalities, input and output data types of tools. The PROPHETS (Process Realisation and Optimisation Platform using Human-readable Expression of Temporal-logic Synthesis) ((24)) program generates suitable candidates of workflows which match the goal of the proposed workflow and its annotations. WINGS offers multiple variations of a workflow created using different tools. It makes use of the input parameters, types of datasets and functions of tools to build the variations ((34)). The approach used by ((13)) utilises data types to facilitate the automatic creation of workflows. All these approaches depend either on annotations or matching input and output data types of adjacent tools in workflows and they pose challenges such as the addition and maintenance of the meaningful annotations of tools and extracting input and output data types of adjacent tools. Moreover, these approaches have their workflow generation restricted to a few specific bioinformatics analyses such as proteomics or proteogenomics. In addition, they do not discuss the presence of higher-order relationships ((22)) in tool sequences of workflows. Our approach to recommend tools in workflows aims to overcome these challenges in the following manner. First, it does not require storing the metadata of tools. Second, it takes into account the higher-order relationships among tools in the tool sequences. Finally, it incorporates workflows from multiple scientific analyses to train the neural network.

### C. Sequential learning on workflows

Workflows, created by many researchers in Galaxy for different scientific analyses, are decomposed into numerous tool sequences (figure 1). The sequential nature of these tool sequences where tools are connected one after another inspires us to apply similar learning techniques used for other sequential data such as text and speech. There are multiple studies in the fields of natural language processing, clinical research and speech recognition which apply deep learning techniques on sequential data to obtain good accuracy in predicting future items. ((36)) finds context in the long sequences of words for sentiment analysis and part-of-speech tagging using RNN and achieves 85% and 93% accuracy, respectively. For clinical data as well, learning on long sequences of health states proves to be beneficial. The health states of patients recorded at different time points are analysed by accessing their electronic health records. The future health states of patients are predicted by training RNN on the sequences of their health states in the past to achieve 85% accuracy ((21)). Moreover, the variants of RNN are used to model speech and music signals ((9); (5)). These successful studies benefit from the sequential learning techniques using different variants of RNN. Therefore, in our work as well, a variant of RNN (GRU) is used to create the tool recommender system in Galaxy. A Bayesian network can also be used for modeling directed acyclic graphs (workflows) ((18); (33)). It requires computing joint and conditional probabilities of nodes in graphs and an increase in the number of nodes can lead to a higher cost to compute these probabilities. In addition, making predictions by learning a probabilistic network is a hard problem ((6);(11);(7)). Because of these drawbacks of using a Bayesian network it is not used in our approach to create the recommender system in Galaxy.

**Fig. 1.**
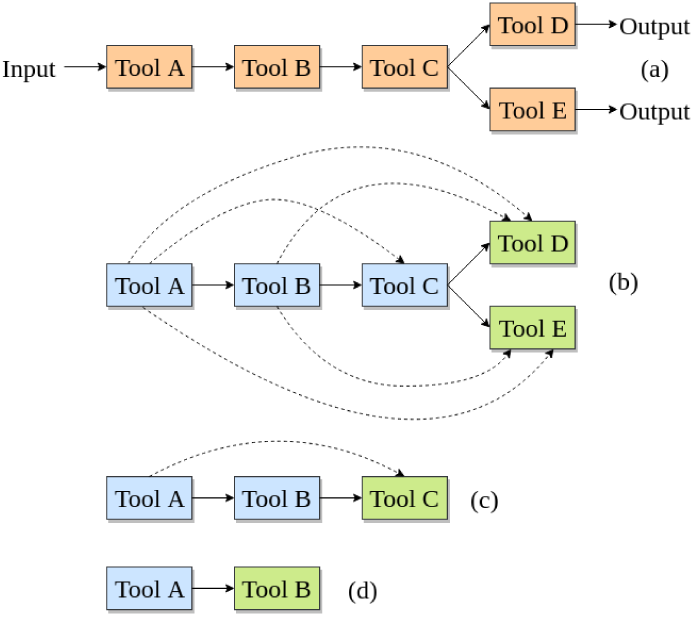
An example workflow (a) is shown consisting of 5 different tools which is decomposed into multiple tool sequences shown in (b), (c) and (d). Each tool sequence shows higher-order dependencies where a tool is dependent on all of its prior tools. These dependencies are shown by the dashed arrows.

## Materials and methods

To create a tool recommender system in Galaxy all the workflows are collected from the European Galaxy server. A workflow may have one or many tool sequences where tools are connected one after another. Tool sequences are transformed into matrices and produced as input to a GRU neural network to learn patterns in the connections of tools.

### D. Data preparation

A workflow (figure 1a) is divided into smaller tool sequences (figure 1b, 1c and 1d). The last tool, shown in green, of each tool sequence (of length n) is assigned as the label of the sub-sequence (of length n-1) shown in blue (figure 1). A label is an output which is learned and predicted by the recommender system. In the neural network learning, a tool is a label. For example, in figure 1b, Tools D and E are the labels of the sub-sequence Tool A → Tool B → Tool C. They show higher-order dependencies in their connections which implies that a tool is not only dependent on its immediate predecessor but also on all prior tools in the tool sequence. For example, in figure 1c, the Tool C is dependent on Tools B and A. By analysing multiple workflow fragments in this way the neural network should learn that the label of a tool sequence Tool A → Tool B is Tool C. It is expected that dividing a tool sequence into fragments with a minimum length of two tools, as shown in figure 1c and 1d, will improve the generalisation performance of the neural network because it gets more tool sequences with a variety of lengths to learn from. The dependencies shown in figure 1b, 1c and 1d present in tool sequences are learned using the GRU neural network by modeling the conditional probability given by equation 1 ((17)). The probability of a tool (*x*_*T*_) is estimated given all other prior tools (*x*_1_, …, *x*_*T*−1_) for a tool sequence (*x*_1_, …, *x*_*T*−1_, *x*_*T*_). The neural network learning is classification because there are labels for tool sequences which are learned and then predicted. Moreover, the classification is multi-class (multiple tools as labels) and multi-label (multiple tools as labels for a tool sequence) ((35)). To ensure an unbiased learning and evaluation by the neural network, the set of tool sequences is divided into two parts - training and test. The training data is used for learning a model and the test data is used for evaluating the model. 

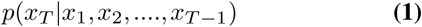

### E. Relevance of tools

The tools in Galaxy have different usage patterns. Some tools are used more often than other tools for multiple reasons such as differences in their functions and availability of similar but better tools. It is essential to analyse the usage patterns of tools because the recommender system proposes tools for researchers and these tools should have high relevance to their analyses. One of the key indicators of relevance of tools can be their high usage frequencies. If a tool has been used often in the recent past, it confirms that the tool is relevant. However, if a tool was used often a few years ago but is being used less often in the last six months then the relevance of that tool has certainly declined. The usage frequencies of tools, shown as labels in figure 1, over the past year are shown in figure 2. To incorporate this usage based relevance of tools in the recommender system, the usage frequencies of all the tools used in the last one year have been collected and are used in the neural network training as the weights (logarithm of usage frequencies) of tools. A tool which has been used often (for example Tool B in figure 2) in the past one year is assigned a higher weight than a tool (for example Tool C in figure 2) which has been used less often in the past one year. When tools are recommended a score is assigned to each tool by the neural network. It is expected that a tool with higher weight gets a higher score and a tool with a lower weight get a lower score. To summarise, the relevance of a tool to be used in a workflow decays if its usage drops over time in Galaxy. Alternatively, the relevance of tools can also be ascertained by counting the occurrence of each tool in all workflows and these occurrences can be used as their weights in the neural network training. It may happen that some tools which were used often in the past to create workflows are not used anymore. Therefore, assigning weights to these tools in the neural network training based on their occurrences in workflows may not be a good indicator of their relevance and overall, may not be optimal.

**Fig. 2.**
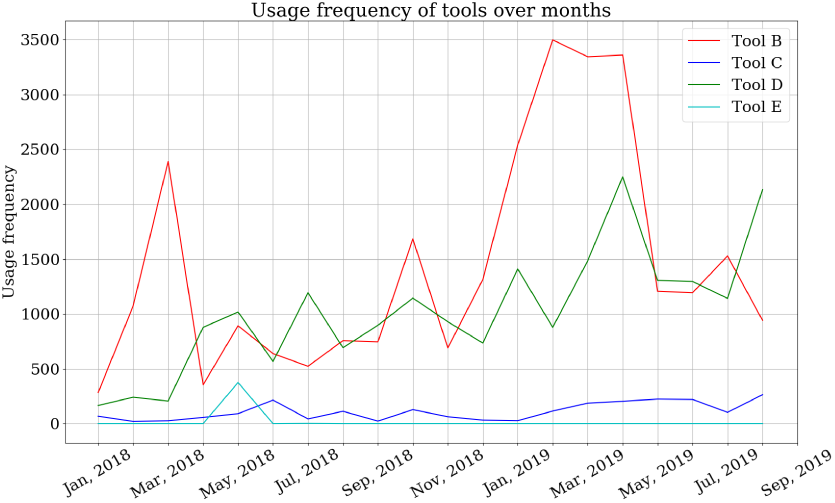
The plot shows the usage frequencies of 4 tools collected over past one year. The Tools B and D have high usage frequencies almost every month while the Tools C and E have much lower usage frequencies compared to Tools B and D.

### F. Implementation

Tool sequences extracted from workflows are transformed into vectors because neural networks require input data to be represented as vectors and matrices. Each tool sequence has one or more labels (figure 1) and they are transformed into different vectors - a tool sequence vector (figure 3b) and a label vector (figure 3d). To form these vectors a dictionary of tools is needed which stores an index for each tool. Using the indices of tools a tool sequence vector is created preserving the original order of tools as in the tool sequence. For example, Tool A has an index of “12” in the dictionary, therefore it is replaced by “12” in the vector (figure 3b). The vector is padded with trailing zeros to keep the length of the vector same across the varying lengths of tool sequences. The size of this vector is 25 which means that a tool sequence can have a maximum of 25 tools. The tool sequences larger than this size are discarded. The labels (figure 3c) are transformed into a bit vector (figure 3d) in which the positions, stored as indices in the dictionary of tools, of the labels (tools) are turned “on” (set to 1) specifying that these tools are the labels of the tool sequence and others are not (set to 0). It has the same size as the dictionary of tools. In machine learning field, it is also known as multi hot-encoded vector. Together, these two vectors form a training sample for the neural network. A pair of vectors are created in this manner for each tool sequence and for all the tool sequences they are combined to form two matrices - one for tool sequences and another for their respective labels. These matrices form input data to the neural network which learns patterns of connections in tool sequences and maps them to their respective labels during training.

**Fig. 3.**
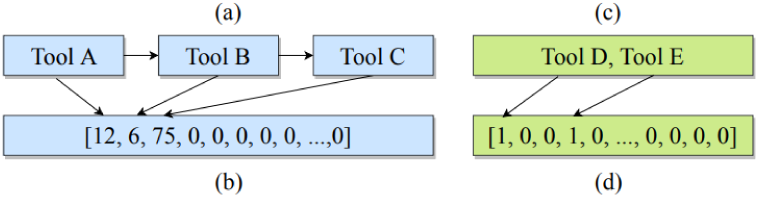
The figure shows how a tool sequence and its labels are transformed into vectors.

#### F.1. Neural network architecture

GRU, a variant of RNN, is used for creating a model which recommends tools. The neural network architecture has four different components (layers) serving different purposes (figure 4).

**Fig. 4.**
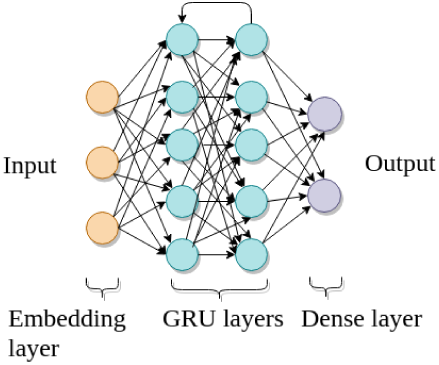
The image shows the architecture of the GRU neural network. It has four components as layers. The first layer is the input layer (yellow), two stacked layers of GRU (cyan) and the last layer is the output layer (violet). The dropout layers are added between two other layers.

### Embedding layer

The first component of the neural network architecture is an input layer (figure 4) which learns an embedding, a fixed-size vector, for each tool. This vector is used by the neural network as an internal representation of a tool. The embedding vector replaces the toolś index in each tool sequence. The size of the embedding vector is fixed for all tools. For example, the vector of a tool sequence [12, 6, 75, 0, 0, …, 0] is transformed into [[0.3, 0.01, 0.003, …, 0.23], [0.5, 0.1, 0.005, …, 0.9], […], 0, 0, …,0] by the embedding layer. The same embedding vector represents a tool in all tool sequences in which the tool is present.

### GRU layer

The stacked layers of GRU learn deeper structures in the tool sequences by modeling the conditional probabilities of tools (labels) given all other prior tools (figure 4). GRU has certain advantages which helps it to learn on sequential data. First, it avoids the problems of vanishing and exploding gradients which commonly occur in traditional RNN ((26)). It is important because learning higher-order dependencies depends on the gradients of errors concerning the parameters (recurrent and input weight matrices) of GRU layers. Second, GRU has slightly fewer parameters than the long short-term memory network (LSTM), another variant of RNN, which makes using GRU simpler than LSTM. Finally, it achieves similar accuracy as the LSTM ((9)).

### Output layer

The last component of the neural network architecture is a dense layer which computes the predictions (figure 4). The dimension of this layer is equal to the number of unique tools because it predicts a score for each tool (label). The predicted score of each tool is considered as its probability of being the label of an input tool sequence. The closer the predicted score of a tool is to 1, the more probable it is to be the recommended tool and the closer it is to 0, the less probable it is to be the recommended tool.

### Dropout layer

Overfitting happens when a neural network performs exceptionally well on the training data but its performance on test (unseen) data remains poor. To prevent it dropout is used between two layers of the neural network. It works by setting some randomly chosen connections in the neural network to 0 ((37); (15)). 3 dropout layers are used in our approach - one between the embedding and the first GRU layers, one between 2 GRU layers and the last one between the second GRU and dense layers.

### Activations

They are mathematical functions which are used in neural networks to transform inputs to a layer into its outputs. Two activations are used in this work - one is exponential linear units (ELU) ((10)) and another is sigmoid (equation 2). ELU is used for both the GRU layers and has a special feature of being negative when the input is negative which allows mean activation (output) to get closer to 0 compared to other activation functions such as ReLU ((23)) which is always positive. As mean activations get closer to 0, the approximated and actual gradients get closer to each other. Therefore, using ELU in our neural network as an activation can be useful to achieve faster training and an increased drop in loss and better accuracy. Sigmoid is used in the output layer which normalises any real number to lie between 0 and 1 and it is considered as a probability of each tool. 

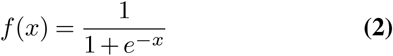

### Usage frequencies of tools as weights

To ensure that the relevance of tools decays with time if they have not been used regularly in the recent past, their usage frequencies are used as their respective weights in the neural network training. The usage frequencies of tools over last 1 year (figure 2) have been collected from Galaxy. A curve is fit through the usage frequencies of each tool using support vector regression (SVR) to display a trend of the toolś usage over time. Using this trend, the usage of the tool for the next month is predicted and its logarithm is used as the weight for this tool. The logarithm of usage frequencies is computed to normalise them because only a few tools have significantly large magnitude of usage compared to that of the remaining tools which may lead the neural network to learn and predict only tools with very large magnitude of usage and ignore other tools. Learning a trend for each tool involves 5-fold cross-validation and optimising two hyperparameters of SVR, kernel and degree, using grid search. The values used for the kernel are - “rbf”, “poly” and “linear” and the values of degree used are 2 and 3. By following the grid search, there are 3 (kernels) x 2 (degrees) = 6 different combinations of hyperparameters to be verified to find the best curve for each tool ((27)).

### Loss function

A neural network learns patterns from data by minimising a loss function. Cross-entropy is a popular choice for a loss function in classification problems ((16)). In our approach cross-entropy function is used in the GRU neural network to compute the loss between the true and predicted label and is weighted by the label’s weight. The loss is summed up over all labels of a tool sequence and then averaged (equation 3). The term *T* is the total number of labels (size of the label bit vector). The term *w*_*i*_ is the weight of the *i*^*th*^ label. The terms *p*^*a*^ and *p*^*b*^ refer to the true and predicted label vectors for a tool sequence, respectively. In general, the loss is large when *p*^*a*^ and *p*^*b*^ are far away from each other which means that the learning by the neural network is not good. If they are close the loss is low and the predictions are better. When an unweighted cross-entropy is used as the loss function for any classification problem ((29)) then it is assumed that all the predictions have the same weight and it does not differentiate between the more and less dominant labels. In our approach when it is used as a loss function in the neural network, then even though the predicted labels are correct they may not necessarily have large weights and thereby maybe less relevant. Therefore, to reduce the possibility of less relevant labels appearing in recommendations, loss is weighted by the weights of labels. It ensures that if a label with a larger weight is misclassified, which means that the true and predicted values are different, then the overall loss is higher. In this way, the wrong classification of labels with a larger weight is penalised more than the wrong classification of labels with a smaller weight. 

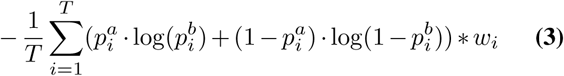

The loss in equation 3 is computed for all tool sequences in training data and is minimised using a root mean square propagation (RMSProp) optimiser. It follows an adaptive approach to estimate the learning rate by keeping knowledge of gradients in prior iterations. The learning rate is updated by dividing it with an average of the square of the prior gradients ((28)).

### Hyperparameter tuning

A neural network has multiple hyperparameters. In our approach they are the number of dimensions of embedding layer, learning and dropout rates, number of units for GRU layer and size of batches. They should be optimised to find the best configuration (a combination of hyperparameters) for training on tool sequences as a different configuration may give a different performance on the same training data. The grid and random searches are popular techniques to optimise hyperparameters. One limitation of these approaches is that they evaluate each configuration independently and have a high time-complexity to find the best configuration. Therefore, the hyperparameters in this work are optimised using a Bayesian (sequential model-based) optimisation ((4)). It learns from the previously evaluated configurations which ensures faster convergence. Reasonable ranges of all the hyperparameters to be optimised are given and the best configuration is found after 30 evaluations.

#### F.2. Learning

The neural network learns patterns in the tool sequences from the training data and creates a model. The ability of the model to recommend tools is evaluated on the test data which is unseen by the neural network during training. While learning, the complete training data is divided into batches of equal size and the weights (belonging to multiple layers of the neural network) are learned in iterations. All these iterations together make an epoch when all the tool sequences in the training data have been used for learning. The number of tool sequences extracted from workflows is approximately 200,000. The training data forms 80% of all tool sequences and it is iterated over 10 epochs of neural network training. The remaining 20% is used as the test data. The running time of the training is approximately 50 hours on Intel(R) Xeon(R) CPU provided by a high performance computing cluster with single core.

#### F.3. Predictions

Learning on training data using a neural network creates a model to predict tools and each tool gets a probability score of being the recommended tool of a tool sequence. The predictions are sorted in the descending order of their probabilities and the top ones (with the highest probabilities) are shown as recommendations. Top-k precision (precision@k) is a popular metric for evaluating a recommender system ((30);(19); (12)). Precision@k implies how many in the *k* predicted tools are correct. For example, *k* = 3 implies that the number of predicted tools are 3 with the highest predicted scores. If only 2 of them are correct, then the precision@3 is 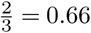. In this way, prediction@3 is computed for all the tool sequences in the test data and then averaged to get an overall precision@3. Precision@1 (top-1), precision@2 (top-2) and precision@3 (top-3) metrics are used in this approach to evaluate the quality of the tool recommender system.

#### F.4. Multiple neural network architectures

Multiple architectures, convolutional neural network (CNN) and dense neural network (DNN) with only dense layers, are used to compare their predictive strengths with GRU neural network (figures 5 and 6). In these architectures too, the embedding layer is used as the first (input) layer and a dense layer is used as an output layer having the same dimensions as the number of tools. Additionally, in CNN, convolutional and max-pooling layers are used to learn spatial patterns in tool sequences and downsample the dimensionality of input, respectively. Moreover, two dense layers are also used and the last one serves as an output layer. DNN uses two dense layers as hidden layers. The cross-entropy, with and without weights, is used as the loss function and RMSProp is used as an optimiser. Bayesian optimisation is used to optimise the parameters these architectures.

**Fig. 5.**
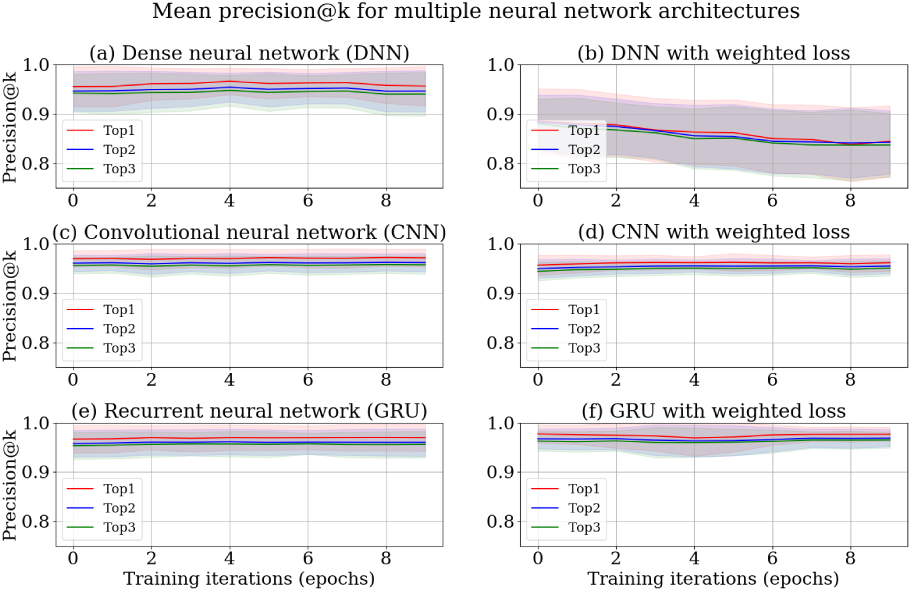
The subplots (a), (c) and (e) show top-k (precision@k) precision for DNN, CNN and GRU neural networks with cross-entropy loss function, respectively. The subplots (b), (d) and (f) show top-k (precision@k) precision for DNN, CNN and GRU neural networks with weighted cross-entropy loss function, respectively.

**Fig. 6.**
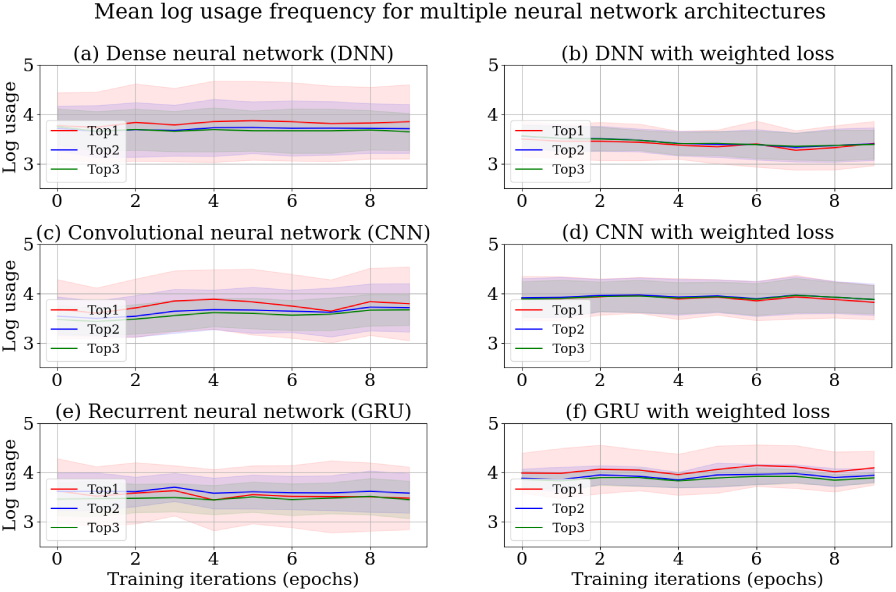
The subplots (a), (c) and (e) show usage frequencies of (top-k) predicted tools for DNN, CNN and GRU neural networks with cross-entropy loss function, respectively. The subplots (b), (d) and (f) show usage frequencies of (top-k) predicted tools for DNN, CNN and GRU neural networks with weighted cross-entropy loss function, respectively.

#### F.5. Library and model

The Keras deep learning library is used for producing the neural network architectures ((8)). The trained model is saved as an H5 file to simplify its distribution to different Galaxy instances. The file is an HDF5 store containing the weights of different layers of the neural network and their configurations, a dictionary of tools and their indices and the weights of tools. The weights and configuration of the neural network are needed to recreate the trained model. The dictionary is used to replace IDs of the predicted tools by their indices in the tool sequence.

## Results

The models obtained after training all the neural network architectures are used to predict tools for the tool sequences in the test data after every training iteration. The precision and usage frequencies of the predicted tools for top-1, top-2 and top-3 metrics are computed over 10 training iterations for each experiment run. They are averaged and their respective standard deviations are computed over 10 experiment runs. The mean precision is shown by line plots and shaded region spans the region between one standard deviation above and below the mean (figures 5 and 6). The GRU neural network with the weighted cross-entropy loss function shows a superior performance to DNN (figure 5a and 5b) by achieving 97% precision (figure 5f) which proves that the GRU layers in a neural network are better for learning on sequential data than the dense layers. Moreover, it shows lower divergence in the means of precision and usage frequencies (figures 5 and 6) establishing that its predictive strength is more stable than DNN over multiple experiment runs. Surprisingly, the weighted cross-entropy loss function does not have any beneficial effect on DNN as its precision deteriorates over training iterations (figure 5b) with a large standard deviation. Due to poor accuracy, DNN is not used in our approach. In contrast to DNN, CNN achieves a similar precision to GRU neural networks with smaller standard deviations (figure 5c and 5d). It also shows an increase in usage frequencies of predicted tools when weighted cross-entropy is used as a loss function (figure 6c and 6d). Despite exhibiting promising results for learning on temporal data (figure 5c and 5d), it gathers lower magnitude of usage frequencies than the GRU neural network with cross-entropy loss function (figure 6d and 6f) which drives it to classify tools with higher usage frequencies more robustly. In other words, GRU neural network with cross-entropy loss function predicts tools with higher usage frequencies and precision than all other approaches. Therefore, it is used in our approach to learn on tool sequences and recommend tools. To illustrate the real-time usage of the recommender system in Galaxy, two examples have been provided - one shows recommended tools for a tool sequence with 3 tools, Trimmomatic → Bowtie2 → FreeBayes in the workflow editor of Galaxy (figure 7) and another displays recommended tools after the execution of RNA-star tool (figure 8).

**Fig. 7.**
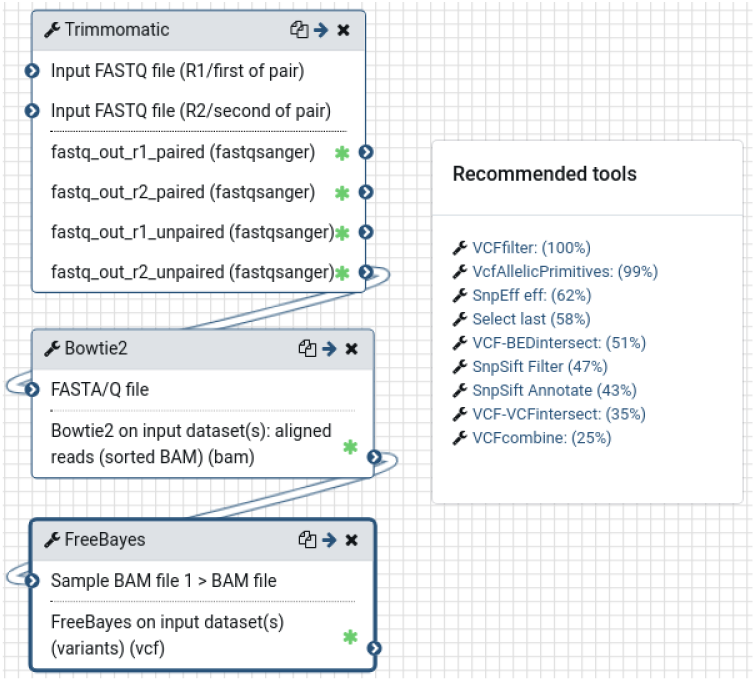
The image shows recommended tools in the workflow editor of Galaxy. The recommended tools can be seen in a modal popup after clicking on the right arrow button placed in top-right corner of each tool. Clicking on any of the recommended tool opens a new block for that tool which can be connected to the tool sequence.

**Fig. 8.**
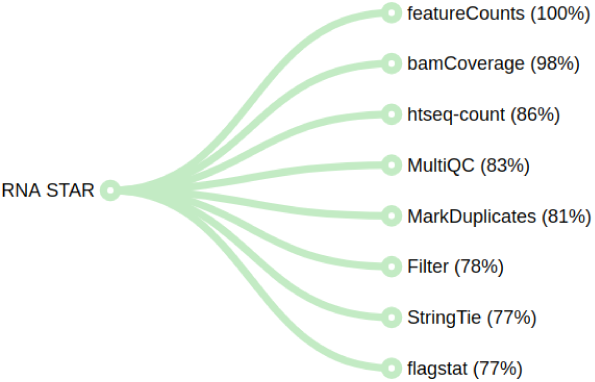
The image shows recommended tools as the leaves (on the right) of the tree after the execution of RNA-star tool. Clicking on any recommended tool opens its definition in Galaxy and can be used for further analysis with the data files produced by the previous tool (RNA-star).

## Discussion

A recommender system to predict tools in Galaxy is built by analysing workflows using a variant of RNN (GRU) and a weighted cross-entropy loss function. The recommended tools are relevant for multiple scientific analyses with a high accuracy, are easily accessible through simple UI integrations and together, they improve user experience by helping researchers to easily create correct workflows. Moreover, the approach does not need to store any metadata of tools and the recommendations are made by only learning the patterns of tool connections in workflows. The model created using this approach is integrated into Galaxy European server to show recommended tools to researchers. An API is provided, residing with other Galaxy APIs, to access a tool or a tool sequence specified by researchers to show its recommendations in real-time. The API is used at two different places in Galaxy - one shows recommendations in the workflow editor and another shows them after each tool execution. The list of recommended tools are sorted in decreasing order of their (predicted) scores. These scores are positive real numbers and are computed independently of one another by the GRU neural network. To make these scores more meaningful, they are normalised by dividing each toolś predicted score by the maximum predicted score.

On a usual Galaxy server the workflows and tools are dynamic, as new tools and workflows are added regularly. Therefore, it is important to train the GRU neural network on the complete set of workflows periodically to keep the tool recommendation model updated with the latest tools and workflows. This model can be created using the set of workflows and tools on a local Galaxy instance by following the steps mentioned in the repository. A script, “extract_data.sh”, is provided for collecting raw data from a Galaxy instance. These are input datasets - one contains workflows and another contains usage frequencies of tools. The sample input datasets are also provided with the scripts. The values of multiple hyperparameters of the neural network, number of training iterations and sizes of training and test data can be altered using the “train.sh” training bash script. To execute the scripts on a GPU enabled machine, the “tensorflow-gpu” package should be installed instead of “tensorflow” as mentioned in the conda package, “environment.yml”, dependencies file. Alternatively, a Galaxy tool is also available to create this model which can be executed directly on Galaxy. This simplifies the creation of a model by providing a UI where the parameters pertaining to the datasets and GRU neural network can be changed. To see recommended tools an ipython script, “tool_recommendation.ipynb”, is also provided which predicts tools for a tool or a tool sequence. A Galaxy admin can overwrite the recommended tools predicted using the trained model by a different set of tools using the Galaxy API which can be beneficial to highlight newly added tools.

## Acknowledgements

We thank Simon Bray for proofreading the paper and Wolfgang Maier for providing feedback. We thank Helena Rasche for providing the data and deploying it on Galaxy European server. The bioRxiv template used here has been adapted from Ricardo Henriques.

## Supplementary information

The project files and workflows are available online. In addition, a tool is created in Galaxy to create the recommendation model.

## References

1. Afgan, E. et al. (2018) The Galaxy platform for accessible, reproducible and collaborative biomedical analyses: 2018 update, Nucleic Acids Research, 46(W1):W537–W544.

2. Baichoo, S. et al. (2018), Developing reproducible bioinformatics analysis workflows for heterogeneous computing environments to support African genomics, BMC Bioinformatics 19, Article number 457.

3. Bela, G. et al. (2009) Scienstein: A Research Paper Recommender System, Conference Proceedings.

4. Bergstra, J. et al. (2013) Hyperopt: A Python Library for Optimizing the Hyperparameters of Machine Learning Algorithms, 12th PYTHON IN SCIENCE CONF. (SCIPY 2013), 2013.

5. Boulanger-Lewandowski, N. et al. (2012) Modeling Temporal Dependencies in High-Dimensional Sequences: Application to Polyphonic Music Generation and Transcription, ICML, 2012.

6. Chickering, D. M. (1996) Learning Bayesian Networks is NP-Complete, Lecture Notes in Statistics, Springer, volume 112, pp 121–130, 1996.

7. Chickering, D. M. et al. (2004) Large-Sample Learning of Bayesian Networks is NP-Hard, Journal of Machine Learning Research, volume 5, pp. 1287–1330, 2004.

8. Chollet, F. et al., Keras, 2015.

9. Chung, J. et al. (2014) Empirical Evaluation of Gated Recurrent Neural Networks on Sequence Modeling, CoRR, 2014.

10. Clevert, D. et al. (2015) Fast and accurate deep network learning by exponential linear units (elus), ICLR 2016, 2015.

11. Cooper, G. F. (1990) The computational complexity of probabilistic inference using bayesian belief networks, Artificial Intelligence, Volume 42, Issues 2–3, March 1990, Pages 393–405, 1990.

12. Deshpande, M. and Karypis, G. (2004) Item-Based Top-N recommender Algorithms, ACM Transactions on Information Systems, Volume 22, Issue 1, pp. 143–177, 2004.

13. DiBernardo, M. et al. (2014) Semi-automatic web service composition for the life sciences using the biomoby semantic web framework, Journal of Biomedical Informatics, Volume 41, pp. 837–847, 2014.

14. Ewels, P. et al. (2016) Cluster Flow: A user-friendly bioinformatics workflow tool. F1000Res, 2016;5:2824.

15. Gal, Y. and Ghahramani, Z. (2016) A Theoretically Grounded Application of Dropout in Recurrent Neural Networks, Proceedings of the 30th International Conference on Neural Information Processing Systems, pp. 1027–1035, 2016.

16. Janocha, K. and Czarnecki, W. et al. (2017) On Loss Functions for Deep Neural Networks in Classification, ArXiv, 2017.

17. Jian, X. et al. (2016) Representing higher-order dependencies in networks, Science Advances, Volume 2, number 5, 2016.

18. Karan, S. and Zola, J. (2016) Exact structure learning of Bayesian networks by optimal path extension, 2016 IEEE International Conference on Big Data (Big Data), pp. 48–55, 2016.

19. Kang, Z. et al. (2016) Top-N Recommender System via Matrix Completion, Proceedings of the Thirtieth AAAI Conference on Artificial Intelligence (AAAI-16), 2016.

20. Leipzig, J. (2017) A review of bioinformatic pipeline frameworks, Brief Bioinform, 18(3): 530–536.

21. Lipton, Z. C. et al. (2015) Learning to diagnose with LSTM recurrent neural networks, CoRR, 2015.

22. Michalski, V. et al. (2014) Modeling sequential data using higher-order relational features and predictive training, CoRR, 2014.

23. Nair, V. and Hinton, G. E. (2010) Rectified Linear Units Improve Restricted Boltzmann Machines, Proceedings of the 27th International Conference on International Conference on Machine Learning, pp. 807–814, 2010.

24. Naujokat, S. et al. (2012) Loose Programming with PROPHETS, Fundamental Approaches to Software Engineering, Springer Berlin Heidelberg (2012): 94–98.

25. Palmblad, M. et al. (2019) Automated workflow composition in mass spectrometry-based proteomics, Bioinformatics, Volume 35, 4 (2019): 656–66.

26. Pascanu, R. et al. (2012) Understanding the exploding gradient problem, ArXiv, 2012.

27. Pedregosa, F. et al. (2011) Scikit-learn: Machine Learning in Python, Journal of Machine Learning Research, vol. 12, pp. 2825–2830, 2011.

28. Ruder, S. (2016) An overview of gradient descent optimization algorithms, ArXiv, 2016.

29. Sadowski, P. (2016) Notes on Backpropagation, Department of Computer ScienceUniversity of California Irvine, 2016.

30. Said, A. et al. (2013) A Top-N Recommender System Evaluation Protocol Inspired by Deployed Systems, 2013.

31. SGomez-Uribe, C.A. et al. (2016), The Netflix Recommender System: Algorithms, Business Value, and Innovation, ACM Transactions on Management Information Systems TMIS, Volume 6 Issue 4, Article No. 13.

32. Smith, B. and Linden, G. et al. (2017) Two Decades of Recommender Systems at Amazon.com, IEEE Internet Computing, Volume 21 Issue 3, Pages 12–18.

33. Spirtes, P. et al. (2000) Constructing Bayesian Network Models of Gene Expression Networks from Microarray Data, Research Showcase @ CMU, 2000.

34. Srivastava, A. et al. (2018), Semantic workflows for benchmark challenges: Enhancing comparability, reusability and reproducibility, PSB (2018).

35. Tsoumakas, G. and Katakis, I. (2007) Multi-label classification: An overview, International Journal of Data Warehousing and Mining, pp. 1–13, 2007.

36. Yin, W. et al. (2017) Comparative Study of CNN and RNN for Natural Language Processing, ArXiv, 2017.

37. Zaremba, W. et al. (2014) Recurrent Neural Network Regularization, ArXiv, 2014.

